# tbea: tools for pre- and post-processing in Bayesian evolutionary analyses

**DOI:** 10.1101/2024.06.18.599561

**Authors:** Gustavo A. Ballen, Sandra Reinales

## Abstract

Estimating phylogenies in which branch lengths are expressed in units of absolute time is crucial for testing hypotheses in evolutionary biology. However, bioinformatic tools to pre- and post-process data from Bayesian divergence time estimation analyses are often not easily interoperable, and documenting methodological choices is not a generalized practice. The R package tbea is a tool-set to integrate biological, geological and palaeontological information to optimize the specification of models, their parameters and prior distributions in divergence times estimation analyses. tbea implements statistical models to (i) better translate time information in dating sources into the specified calibration densities, (ii) improve comparisons between prior and posterior distributions for parameters of interest, (iii) carry out inference on origination times for a set of distributions, (iv) summarise different distributions into a single one, and (v) improve the reproducibility of divergence time estimation analyses allowing users to document methodological choices. We illustrate the package functionalities by carrying out two worked examples. One on the phylogenetic relationships and divergence time estimation of South American Cynodontidae, and another one on the separation time of drainages East and West of the Andes in South America. It is expected that the tools herein available will be key when estimating events in time from sets of point estimates, as well as the combination of different posterior densities from the same parameter are useful to justifying the selection of secondary calibration points, or discussing the timing of biogeographic events when multiple sources are available.

## 1 Introduction

In phylogenetic inference, the characters, either molecular or morphological, only contain information about how closely or distantly related the terminals are. Branch lengths, as estimated in standard phylogenetic analysis (e.g. ML or Bayesian inference) represent a combination of the expected number of changes per site and the time required to accumulate changes along branches. Therefore, estimating a phylogeny in which branch lengths are in units of absolute time is key in comparative analyses such as modelling morphological evolution, biogeography, or macroevolutionary dynamics (Warnock and Wright, 2021). One key reason why time trees are useful lies in the nature of evolutionary models which consider a stochastic process in time, such as the discrete Markov process often used to model evolution (Lewis, 2001; Yang, 2014).

In a very general form, we can describe a Bayesian divergence time estimation model as

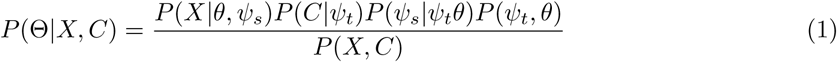

where *P* (Θ|*X, C*) is the posterior distribution of the model parameters, *P* (*X*|*θ, ψ_s_*) is the phylogenetic likelihood of the sequence data *X*, *P* (*C*|*ψ_t_*) are the time calibration densities, *P* (*ψ_s_*|*ψ_t_θ*) is the clock model, *P* (*ψ_t_, θ*) is the tree process, and *P* (*X, C*) is a normalising constant.

The denominator *P* (*X, C*) in Equation 1 cannot be calculated analytically and we need resort to MCMC sampling algorithms for approximating the posterior distribution(s) numerically. The result is a posterior sample of time trees in which branch lengths are in units of absolute time (usually million years, Ma). Node times and branch lengths are parameters in the tree topology *ψ*, and the posterior age distribution of each node can be summarised from the sample of time trees. When the tree *ψ* is fixed, we only have the posterior distributions of node ages, but when the tree is coestimated with the divergence times, we also have a posterior distribution of trees, which may or may not share nodes across the different elements of the distribution. A Bayesian analysis of any sort aims to characterise the posterior distributions and thus the uncertainty on estimated parameters rather than point estimates. Terminology applied to divergence time estimation has been quite inconsistent in the literature.

It has been called molecular clock analysis, tree calibration, molecular dating, tree dating and clock dating in the massive literature on theoretical and empirical aspects of this family of analyses. It is useful to define here the different terms in use as these allow a more precise description of the elements in the analysis. We propose to use divergence time estimation as the name for the analysis where branch lengths are separated in rate and time and results in a time tree where branch lengths represent time. Clock, be it molecular or morphological (Barba-Montoya et al., 2021), is a model that describes how the rates evolve along the tree. Calibration is the specification of time information in the form of a calibration density, which is a probability density function that describes the uncertainty of nodes or tips times in the tree. Dating means telling the age of an object, for instance with the use of geochronological, biostratigraphic, or any other information which allows to tell the absolute age of fossils. In this sense, none of trees, clocks, or molecules are dated but instead the fossils which are used to construct calibrations. Note that dating is a term widely applied to geochronological research (Rink and Thompson, 2015). Under this scheme, molecular (or morphological) clocks as well as calibration densities are components of the complete divergence time estimation model depicted in Equation 1. We use the terms and definitions above throughout the present article.

The most common strategy for using independent temporal evidence to calibrate a phylogeny to absolute time is assigning time constraints to one or more nodes in the phylogeny, usually called node calibrations (Rannala and Yang, 2007). This time information comes from a variety of sources, including geochronological estimates from fossils or biogeographic events (primary calibrations; Landis, 2017) or time estimates obtained from previous studies (secondary calibrations; Schenk, 2016). Node calibrations based on fossil evidence are set using the oldest fossil record of a clade, which establishes a minimum age for that node in the phylogeny (Rannala and Yang, 2007; O’Reilly et al., 2015). Nevertheless, it is possible to consider the fossils are part of the same diversification process that generated the extant species, and incorporate them as dated tips alongside living relatives (tip calibration; Ronquist et al., 2012; Heath et al., 2014). This approach allows for inclusion of all available fossils (Heath et al., 2014), but the impact of the taxonomic and the stratigraphic age uncertainty propagates to divergence time estimates and should be incorporated explicitly (O’Reilly et al., 2015; Barido-Sottani et al., 2019). Several methods are available to estimate time trees using one of these approaches (e.g.; To et al., 2015; Sanderson, 2002; Stadler, 2010; Heled and Drummond, 2009).

One of the challenges in node calibration strategies is translating the inherent uncertainty in fossil age information into a prior distribution (Warnock et al., 2015). This uncertainty usually comes from the litho-, bio-, chemo-, cyclo-, and /or magneto-stratigraphic method using to establish the absolute date of the fossil provenance (O’Reilly et al., 2015; Parham et al., 2012). There are a variety of probability densities implemented to account for the fossil age uncertainty (e.g., Uniform, Lognormal Warnock et al., 2015). Some programs such as Beast2 (Bouckaert et al., 2019) require users to input parameter values for the node calibration densities, often by eye, aided by plots as is done in beauti. This process is not reproducible and can be impractical when several calibration points must be set.

On the other hand, when different time estimates for the same event (e.g., a node) are available, summarising such information to constrain a node age (i.e., secondary node calibration) using all available information is not straightforward. Consequently, empirical studies usually resort to manual specification to find a single and representative distribution, or choose among alternatives using all sorts of arguments such as the most cited estimate for the group of interest, the most recent, or the one with the best taxonomic sampling. Nevertheless, a reproducible method to summarise different sources of information would allow incorporating the uncertainty associated with the time estimates for a node, because every study constitutes evidence, as long as the estimates are independent. Tools for summarising different time estimates can be also applied in biogeography. Empirical approaches to dating vicariant events in specific groups involve plotting the time estimates along with their uncertainties to visually extract information on when such events are likely to have occurred (Hazzi et al., 2018; Pérez-Escobar et al., 2019). Another alternative has been to plot the mean of these data as a point estimates for the general time of separation (Smith et al., 2014). Nevertheless, a statistical approach would allow to infer the age and its uncertainty, at which the process responsible for all those vicariance events took place (e.g., separation time of drainages East and West of the Andes in South America), using divergence time estimations for different biotic groups. These tools will impact beyond evolutionary biology as Bayesian phylogenetic methods are also applied to other fields in cultural evolution such as linguistics (Maurits and Griffiths, 2014; Hoffmann et al., 2021; Neureiter et al., 2022; Greenhill, 2023), palaeolithic artifacts (Matzig et al., 2024), and musicology (Hajic Jr et al., 2023; Ballen et al., 2024; Hajic jr et al., 2025).

The package tbea has tools for improving the reproducibility of the generation of concatenated alignments *X*, express better the time information in calibration sources onto the specified calibrations *P* (*C*|*ψ_t_*), propose adequate initial values for the time tree *ψ_t_*, improve comparisons between prior and posterior distributions for parameters of interest, in particular node times, and carry out inference on origination times for a set of posterior distributions of the same class in different studies. The package also includes tools for summarising tree distributions in terms of both topology and branch lengths, as well as two methods for estimating the time of an event where independent estimates (*P_i_*_:*N*_ (Θ*_i_*|*X_i_, C_i_*)) are available (e.g. multiple estimates of the timing of a biogeographic event). Finally, it implements conflation of densities as a tool for summarising multiple distributions of the same parameter into a single one. This tool-set will help researchers in evolutionary biology to improve the reproducibility of their Bayesian analyses, and help them streamline coding strategies in different phases of analysis, from specification to comparisons of results.

Typical users of tbea include evolutionary biologists wanting to use Bayesian methods in the realm of phylogenetics and macroevolution where it is necessary to specify temporal information in the form of calibration priors. Researchers in palaeontology and geology may also use functions of this package when interested in format conversion of morphological matrices, summarisation of phylogenetic analyses of extinct and/or extant organism, combination of multiple time estimates in the form of distributions, and event tools for testing the assumption of isochron. Finally, any researcher interested in combining multiple distributions may find the implementation of distribution conflation helpful, even outside the realm of biology as it is a general statistical technique. Finally, researchers applying statistical methods developed in a biological context but later applied to questions in cultural evolution and digital humanities may use the tools herein presented in their own research questions.

## 2 Implementation

Three computational techniques are used in several functions of this package: Quadratic loss optimisation, numerical integration, and metaprogramming. Optimisation is used when fitting parameters of interest, as well as during calculation of quantiles. This is achieved by using numerical optimisation on a quadratic loss function of the form argmin*_x_ ϕ*(*x*) = (*f* (*x*) − *t*)^2^ where *f* (*x*) is the target function and *t* the target value. Optimisation stops once |(*f* (*x*) − *t*)^2^| *< δ* is attained for a given tolerance level *δ*. Optimisation works with both univariate and multivariate functions. This technique is used in the functions findParams, c truncauchy, and quantile conflation.

Numerical integration uses an approximation for the area under the curve 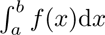 where *a* and *b* can be positive and/or negative infinite as well as real numbers. It approximates the area under the curve using adaptive quadrature in unidimensional functions where multiple subdivisions of such area are approximated and proposed following an adaptive procedure that decides when to stop subdividing the area, once *ɛ < δ* where *ɛ* is the error and *δ* is the tolerance, in a similar way as in numerical optimisation. This is used for calculating quantiles on arbitrary conflated distributions, as well as for calculating the conflation itself (see below).

Metaprogramming works by writing code that returns code which is in turn evaluated. For instance, suppose a function returns a string which constitutes a valid expression expr = ‘1 + 1’. Then, the value of expr can be parsed and evaluated by the interpreter as its input: eval(expr) => eval(‘1 + 1’) => 2. The power of such technique is that the user does not need to hard-code expressions for calculating operations that take functions and their evaluations as input, and therefore the output of such evaluations is very flexible. This is especially useful for implementing operations that take functions as input rather than concrete values such as vectors, matrices, or lists. This technique is used when generating function calls using probability density functions in findParams as well as for conflating distributions.

The package tbea is hosted on CRAN (https://cran.r-project.org/package=tbea) and development versions are available on GitHub (https://github.com/gaballench/tbea). The stable version of the package can be installed with library(“tbea”) and the development version with remotes::install github(“gaballench/tbea”).

### 2.1 Dependencies

One of our design choices was to reduce dependencies to a minimum, which at the moment depends on ape (Paradis and Schliep, 2019) for phylogenetic tree representation and manipulation, Rfit (Kloke and McKean, 2012) for extending the possible linear models to be used when inferring the *x*-intercept of cumulative density functions, and coda (Plummer et al., 2006) for calculating the highest posterior density interval which is used in crossplots.

### 2.2 Comparison with other packages

Several packages have been developed for phylogenetics and related areas (Gearty et al., 2024). There are as many standards as software tools, and this is reflected in the multiplicity of approaches to software development applied to evolutionary biology. Multiple functionalities are also implemented in different ways, thus providing a high amount of flexibility to users of phylogenetic methods. Some tools available in this package are already available in a different form in some other packages (e.g. tools for interacting with data matrices, concatenation, and format conversion), while others are completely new to our knowledge (e.g. parameter estimation in distributions with a set of quantiles, confidence intervals on the x-intercept, or the conflation method). We think that having multiple implementations of the same tools is positive rather than redundant as much as the different implementations allow the same tools to be used in different analytical contexts rather than forcing the users to have the same settings, formats, and input.

Some packages target Beast or Beast2 as the preparation of input, programmatic use, and parsing of output is challenging under certain circumstances. One of the most popular tools, beauti (Bouckaert et al., 2019), is used as a companion for Beast and Beast2 as it allows to generate the XML input file from a graphical user interface. The package beastier (formerly babette; Bilderbeek and Etienne, 2008) on the other hand, has code for interacting with programs of the Beast suite. Although some of our functions can be used to help specify analyses in Beast2 and parse output logs from the program, our package is not intended to be a wrapper nor a gateway to Beast, but rather a tool for improving reproducibility when using that program. A similar package, beastio (du Plessis, 2020), aims at providing pre- and post-processing tools specifically for Beast and Beast2.

EvoPhylo (Simõoes et al., 2023) targets divergence time estimation analyses using morphological datasets and their clock models, although it can be used for molecular datasets too. The package also pre- and post-process output from both Beast2 and MrBayes but restricted to situations where morphological data, in particular for (but not restricted to) completely extinct taxon sampling, is present in the analysis. In a similar way, Chronospace (Mongiardino Koch and Milla Carmona, 2024) targets the exploration of time-tree parameter distributions and measures of sensitivity when carrying out multiple analyses. Our package has functionality which similarly targets unrooted tree distributions, thus being complementary. Convenience (Fabreti and Höhna, 2022) is a package for assessing Bayesian convergence both in continuous and discrete (topology) parameters, therefore offering post-analysis tools. RevGadgets (Tribble et al., 2022) is tightly integrated with RevBayes in a similar way as packages targeting Beast/Beast2. It has functionality for assessing estimated distributions similar to Tracer but implemented in R, and therefore offers tools similar to our tools for comparing distributions. It also has tools for plotting trees from a number of analyses, thus offering output similar to FigTree (Rambaut, 2018) but using R code. It is strongly integrated with ggplot2 whereas our approach is to use base instead in order to reduce the amount of dependencies while producing simple yet informative plots. CladeDate (Claramunt, 2022) is very similar in that it aims at generating calibration densities from the fossil record. However, that package uses data argumentation via sampling from the uncertainty in fossil occurrences in order to generate a distribution of a parameter *θ* (fitted by using also methods for stratigraphic intervals) representing the origination of the clade. phruta (Romàn-Palacios, 2023) has tools for retrieving data from GenBank and then concatenate them for phylogenetic analysis. In contrast, the present package allows to concatenate matrices including both DNA and morphological data, so that total-evidence divergence time estimation and tree inference can be carried out combining these two data sources. The package mcmc3r (dos Reis et al., 2018; Álvarez-Carretero et al., 2019) targets analysis such as model selection using Bayes factors specifically using the program MCMCTree, whereas tools in tbea can be used for comparing results form independent runs using MCMCTree. Also, mcmc3r has functions for plotting the minimum and both upper and lower bound calibration densities used by MCMCTree, in a similar way as our package aids in plotting Beast2’s Lognormal calibration density. Finally, TreeTools (Smith, 2019) implements tools for interacting with output from TNT (Goloboff et al., 2008). Although tbea has a function for reading trees in TNT format, its aim is to convert them to newick format rather than to generate an R representation of the tree; therefore, the operations on the TNT tree text representation return also a tree in text representation, but in newick format instead.

### 2.3 Worked examples

We introduce the package and illustrate its functionalities by carrying out two worked examples. One on the phylogenetic relationships and divergence time estimation of South American Sabre-Tooth Characins of the family Cynodontidae, and another one on the separation time of drainages East and West of the Andes in South America. The former targets pre-analysis tools and a couple of functionalities for post-analysis settings, whereas the latter aims at demonstrating the use of post-analysis tools when information from different analysis is available and the goal is to infer a single parameter, in this case, the separation time between these two biogeographic regions.

#### 2.3.1 The South American Sabretooth Characins

We will use the South American Sabre-Tooth Characins of the family Cynodontidae for a divergence time estimation analysis. The phylogenetic relationships of the group were revisited by Ballen et al. (2022), although these authors did not provide information about the temporal scale of their evolutionary relationships. We will use the morphological and molecular datasets of that study, including a fossil terminal.

The dataset comprises 10 terminals, eight molecular markers, and one morphological partition with 80 characters. We refer the reader to the study for details on the substitution models and other details of the phylogenetic analysis.

Divergence time estimation was carried out using Beast2 (Bouckaert et al., 2019) with the SampledAncestors package (Gavryushkina et al., 2014). An analysis sampling from the prior, that is, ignoring the alignment, was also carried out with the option sampleFromPrior in beauti. Preparation of the initial tree was carried out using the package ape (Paradis and Schliep, 2019). Post-analysis tree plotting was carried out in R (R Core Team, 2024).

#### 2.3.2 The separation of drainages East and West of the Andes

Divergence time estimation data were compiled from the literature that reported separation events between cis- and trans-Andean lineages, be it contemporary species or nodal divergences (Abe et al., 2014; Cheviron et al., 2005; Collins and Dubach, 2000; Cortés-Ortiz et al., 2003; D’Horta et al., 2013; Dick et al., 2003; Elias et al., 2009; Fernandes et al., 2014; Grau et al., 2005; Gutiérrez et al., 2014; Hardman and Lundberg, 2006; Herńández Torres, 2015; Machado et al., 2014; Miller et al., 2008; Patané et al., 2009; Patel et al., 2011; Picq et al., 2014; Ribas et al., 2005, 2007; Říčan et al., 2013; Ruiz-García et al., 2015; Smith et al., 2014; Voss et al., 2013; Weir and Price, 2011). A total of 54 data points were compiled along with their credible intervals, most frequently highest posterior density intervals.

The distribution of divergence time data is very asymmetrical with a heavy right tail, product of several outliers older than 10 Ma (Figure 1). These outliers affect strongly those methods who make a stronger assumption on the lack of uncertainty in the measurement of time values (e.g., methods based on stratigraphic intervals and less strongly, those based on cumulative densities).

**Figure 1:**
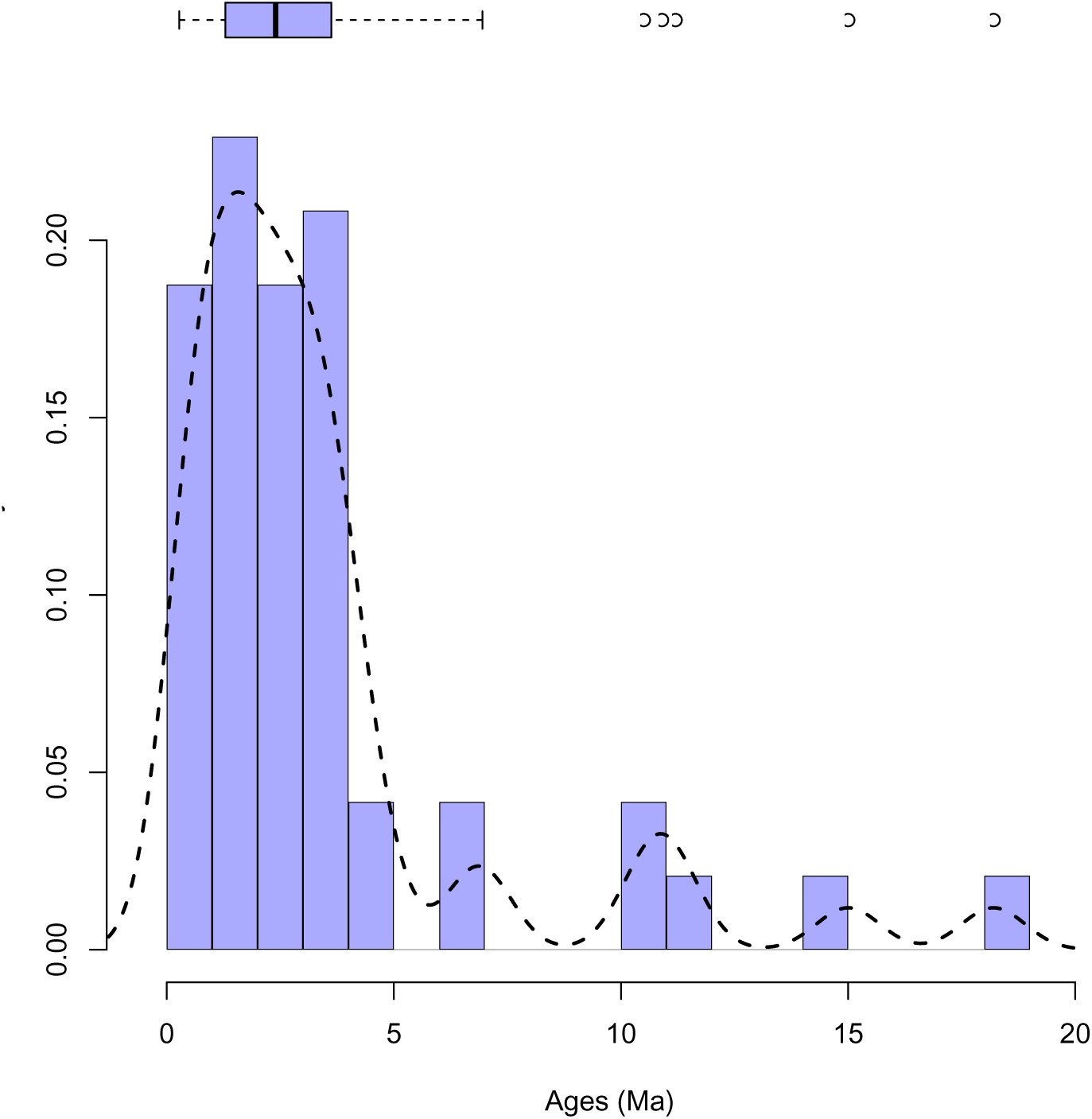
Descriptive statistics of the divergence time data. The histogram illustrates that the multimodality in the density line is an artefact of discontinuity in the age data, while the boxplot on top of the figure shows the existence of multiple outliers to the right of the distribution. All three methods agree in showing the asymmetry in the distribution with a heavy right tail. Discontinuous line represents the empirical probability density.

#### 2.3.3 Coding examples and vignettes

The package tbea provides a number of vignettes in addition to the examples in this article. PDF rendered versions of the vignettes can be accessed both on CRAN (https://CRAN.R-project.org/ package=tbea) as well as on a dedicated website https://gaballench.github.io/tbea.

## 3 Results

### 3.1 Concatenation and total-evidence analysis

Concatenation can mean two different operations when working with partitions of data in phylogenetic analysis. On one hand, concatenation means pasting together multiple data matrices (i.e., data partitions) of the same data type (e.g., DNA alignments) into a single one which is in turn analysed as a single partition, with a single evolutionary model and therefore parameter set. There are numerous issues when combining data partitions in this way, with the fact that different genome regions resulting in different gene trees probably the most problematic one. As a consequence, the largest original partition will end up dominating the signal during inference because the parameter set is estimated from the partition as a whole. On the other hand, concatenation means combining different partitions into the same analysis and informing the same tree, although applying independent and potentially different evolutionary models for each. Independent partitions can now inform equally shared parameters (i.e., branch lengths and topology), while having their independent parameter sets estimated independent from one another. Because of this property, it allows to use multiple data types as partitions in what has been called total-evidence analyses (Ronquist et al., 2012; Heath et al., 2014), where we may for instance have some partitions for specific genes, a partition for aminoacid data, and yet another one for morphological characters. This form of concatenation does not however solve the issue of gene–tree to species–tree discordance (in fact, it can be still very problematic in these scenarios; Mendes and Hahn, 2018), but is the only alternative available when multiple data types are of interest. In this work we use the second meaning of concatenation as it is the relevant one during divergence time estimation using total evidence approaches.

Tools are available for concatenating DNA matrices (e.g. the R packages apex; Jombart et al., 2017), although few deal with concatenating different data types such as morphological matrices (but see Salinas et al., 2024). Our package offers tools designed for including both DNA and morphological matrices (see the vignette concatenation), which is necessary e.g. in MrBayes for total-evidence analyses. Often the morphological datasets are compiled in some form of table, for instance a spreadsheet, and the conversion to nexus for later concatenation to other partitions is a useful feature even for working with programs that support individual files per partitions such as RevBayes or iqtree.

### 3.2 Calibration specification

Calibration densities can be divided into two groups: Node and tip calibrations. Node calibrations are probability density functions which represent our uncertainty about the age of a given node. Similarly, tip calibrations represent the uncertainty about the age of a terminal in the tree, for instance, a fossil terminal for which we also have data in matrix. Different programmes make available different probability density functions, being the Normal, Lognormal, Exponential, and Uniform popular options. Calibration densities may also represent bounds on an age, that is, lower and upper quantiles of some probability, e.g. the interval of 95% of the probability. In the case of continuous distributions with support in (−∞, ∞), [0, ∞), or (0, ∞), one or both bounds are soft, that is, we allow ages to be outside of that interval with some probability, being it 2.5% on each tail if we use a 95% interval. In other distributions such as the Uniform, bounds can be hard, that is, we say the probability of being outside the interval is 0. It is possible to set distributions with combinations of soft and hard bounds. For instance, the Lognormal distribution with an offset says that the probability of observing ages younger than the offset is 0, whereas the upper bound corresponding to a quantile of 95% will be soft with probability 5%.

We specify a calibration density by picking a given density function and setting its parameter values. Some considerations are important when setting calibration densities, for instance, if we use the uncertainty from the dating of a given fossil as the calibration density, it implies that we mean the fossil age to be also the node age. This may be a strong assumption about the position of the fossil, and then we may want to use the fossil as a lower bound instead (Parham et al., 2012). In this case, we may want the fossil to represent a hard bound, that is, the age of the node of interest cannot be younger than the fossil, and then pick an upper soft bound for an age such that older ages are still possible yet unreliable. Let us suppose we have a node with a fossil age of 50Ma which we want to set as a hard bound, and then previous estimates of the age of the total group under analysis saying that it is probably younger than 100Ma, for instance being this the root age. We may pick a distribution with a hard bound at 50Ma and a soft 95% bound at 100Ma. Because additional considerations need to be made in this second approach, we will assume that the fossil in the following example is on the node.

Regardless of the specific details supporting our choice of bounds, it is important to use proper densities when specifying calibration priors. Improper densities are those whose area under the curve for the complete support does not integrate to 1.0. In contrast, proper densities integrate to 1.0. Improper densities induce issues during MCMC sampling (Yang, 2014) and should be avoided. An example of improper distribution is setting a calibration density as a Uniform with just a minimum hard bound but no maximum bound *U* (*min,* ∞) with support in [*min,* ∞], because 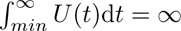.

We will now specify two calibration prior densities to estimate divergence times for the family Cynodontidae: (i) a node calibration using the oldest record of the genus *Hydrolycus* (early Barrancan, *>* 41.6 − 40.94 Ma) to calibrate the most-recent common ancestor (MRCA) of *Hydrolycus armatus* and *H. scomberoides* using a Lognormal distribution, and an alternative representation of the same age uncertainty using an Exponential distribution (see vignette node calibration density). In general, if the uncertainty can be described by either an Exponential or Lognormal, the former is preferred because it has one parameter instead of two in the latter. The difference between both lies in the flexible position for the mean inside the support of the Lognormal distribution, whereas the mean, 0.025, and 0.975 quantiles are governed by a single parameter in the Exponential distribution.

#### 3.2.1 Node calibration density

When using time information associated to a fossil as calibration density, age uncertainty usually come from biostratigraphy (e.g., if no radiometric estimate applies closer in the stratigraphic context to the fossil locality). Then, it is possible to represent that uncertainty using a Uniform distribution with first and last time parameters (e.g., 40.94 and 41.6 Ma respectively), or a Lognormal distribution with soft bounds. In the second case, we can find the combination of mean and standard deviation that better describes a distribution whose density cumulative probability values 0.0, 0.5, and 0.975 correspond to the quantiles defined by the Uniform prior above. The function findParams finds such combination of parameters for a given pair of quantiles. As the standard Lognormal distribution is defined between (0, ∞), we need to apply an offset towards the minimum age. This will make it possible to relax the rather strong assumption about the node age, because by using a Uniform prior the probability of observing times with fall outside the interval is zero.

Note that the Exponential distribution can be parametrized in terms of the scale (1*/λ*, as is the case for Beast2) or the rate (*λ*) as used by R. Therefore the value to input when defining the prior of an Exponential distribution in beauti is, which in our case is 1*/*5.589202 = 5.8. When plotting the prior in R (e.g. using the function curve as above) we need to use *λ* (i.e., 0.17) rather than the mean (Figure 2B).

**Figure 2:**
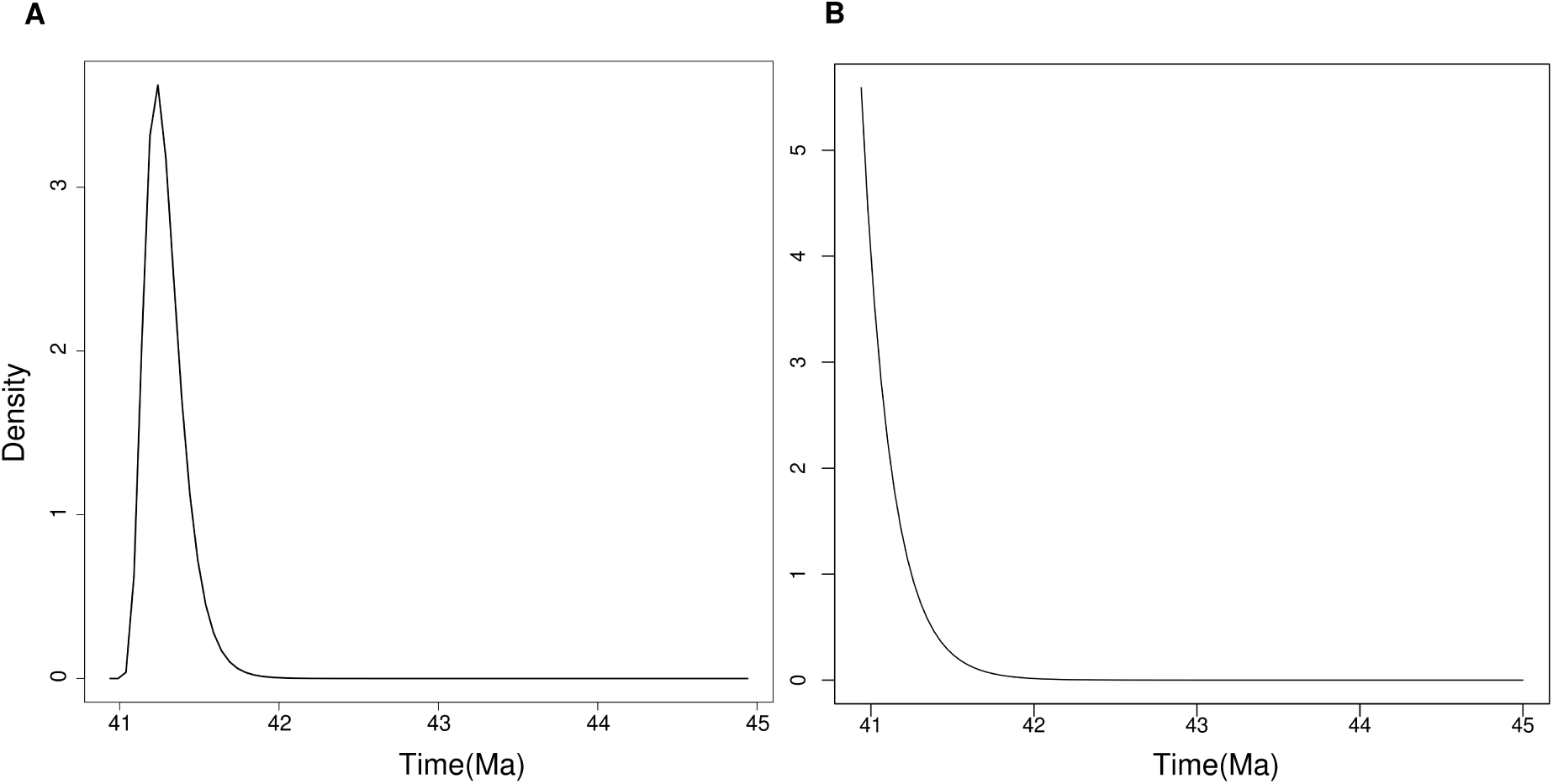
A. Calibration density for the node corresponding to the genus *Hydrolycus* as generated by the function findParams from the quantiles found in the calibration justification and plotted with lognormalBeast. B. Alternative specification of the same calibration density but using an Exponential distribution as generated by the function findParams and plotted with curve(dexp(x)).

#### 3.2.2 Alternative specification: the **L** calibration density of **MCMCTree**

Lets suppose that we want to carry out an analysis using MCMCTree instead (see vignette l calibration mcmctree). That programme uses different densities for specifying a node calibration. One of these, called L implements a truncated Cauchy distribution (Inoue et al., 2009) whose shape can be controlled by the variables location and scale (p and c). It is used frequently for describing minimum-age calibrations where a maximum is unknown. Although it can give the impression that we only need a minimum age (which can be soft or hard), this is not the case of an improper distribution as it is controlled by p and c so that its area under the curve is 1.0. It is useful for describing asymmetric calibrations, although care must be taken as specifying both p and c have a profound impact on the shape of the distribution (see e.g. the unnumbered figure in p. 49 of the PAML documentation; Yang, 2010). The function c truncauchy helps us specify such density as it estimates the value of c provided that we have both age minimum and maximum, and define some probability pr to be allocated to the right of the maximum age. For instance, our minimum age is say 1 in units of 100Ma, and our maximum age is 4.93 also in units of 100Ma. The minimum is a soft bound allowing 0.025 of the density to be allocated to the left of it, whereas 0.975 is the percentile at which we observe the maximum age. The quantity p=0.001 has been chosen so that the mode is closer to the minimum age. Then, we can find the value of c which completes the definition of the L density to be used in MCMCTree, which happens to be around 0.0009.

### 3.3 Initial trees

Programmes that co-estimate the tree topology and divergence times (e.g., Beast2, RevBayes, MrBayes) first need to initialise the tree in order to start MCMC sampling. Depending on the size of the analysis, that is, the number of terminals, such initial tree can be quite bad and make convergence to take longer. It is therefore useful to help initialise the analysis by setting the initial tree, by calculating a reasonable one from which sampling then converges quicker. When doing standard tree inference we do not need to worry about the tree being consistent with times, but this is not the case when doing divergence time estimation. In the latter case, not just the tree but also the node ages need to be consistent with the calibration densities, and we need to guarantee that before starting the analysis. This is a frequent reason for an analysis in Beast2 being unable to initialise.

The package tbea helps us find initial trees when using the output from TNT, a parsimony programme, by providing functions for doing format conversion from TNT to standard newick format. Afterwards, it is possible to use penalised likelihood as implemented in the package ape for finding a set of branch lengths which is consistent with our calibrations as shown in the vignette starting tree. In the Figure 3 we see an example of a tree that was converted from TNT format and then set to branch lengths consistent with the calibrations specified in the previous section.

**Figure 3:**
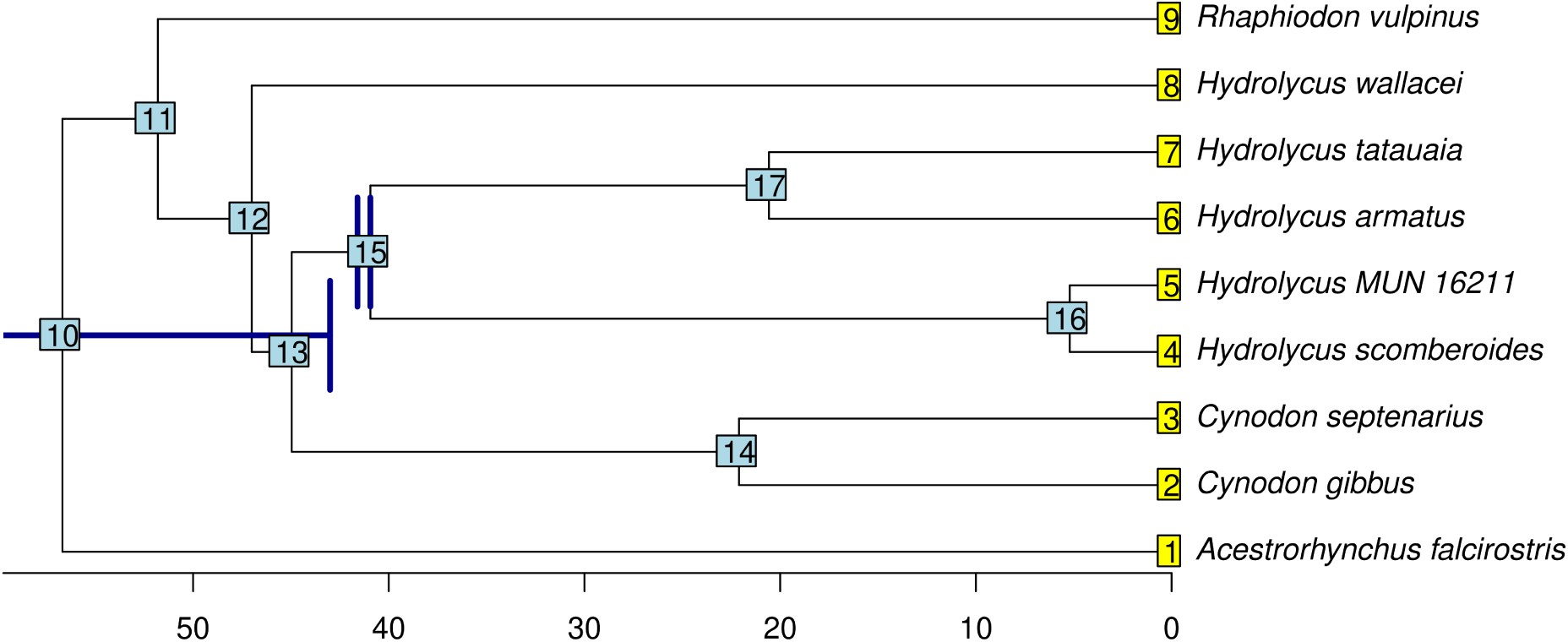
Calibrated tree of the Cynodontidae which is consistent with the calibration densities and can be used as starting tree in analyses of divergence time estimation. Calibrations shown are the root calibration density to the left, and the *Hydrolycus* calibration density to the right.

### 3.4 Post-analysis comparisons

#### 3.4.1 Summary of (posterior) tree distributions

After co-estimating the tree topology during MCMC sampling, we end up with two sets of values, one for parameter values, and another one with a tree sample. Depending on the type of analysis, trees can be unrooted as in standard tree inference, or rooted as in divergence time estimation. Different programmes implement their own summarising tools which fit the different design choices and therefore are not necessarily interoperable: Tools for Beast2 do not necessarily work with samples from MrBayes, and *vice versa*. In particular, existent summary tools for tree topology are not well suited for unrooted trees as they were designed with time trees in mind, which are rooted.

There are two steps necessary for summarising tree samples (see vignette posterior tree distribution). One is to decide how many distinct topologies (disregarding branch lengths) are in the set of trees and then we need to summarise branch lengths for each of these tree subsets using any sensible summary measure (e.g., mean or median). The functions topoFreq and summaryBrlen aid in these two tasks respectively. topoFreq uses the Robinson-Foulds (RF) distance among trees in order to find clusters which are identical in topology and then bind them together in a list that we can later operate over. The RF distance is defined over unrooted trees, so it is necessary to check whether the input trees are. Unrooting a rooted tree will destroy the root node and then add up together both adjacent branch lengths, and this will be reflected when summarising with summaryBrlens using the output of topoFreq. This will return one branch length less than the number present in the original sample of rooted trees. Accordingly, this method will only work on sets of unrooted trees, or sets of time trees of fixed topology where we are only interested in finding a summary values for branch lengths. The function topoFreq should not be used when both the topology and divergence times are being coestimated (e.g., when the posterior of trees comes from not fixing the topology in Beast2, RevBayes, and MrBayes) but it is safe to use when the inference was done to estimate only the unrooted topology and branch lengths, such as in standard tree inference in MrBayes. In the example, we used MrBayes for standard tree inference of the Cynodontidae dataset, and then summarised the posterior tree sample using the topoFreq and summaryBrlen functions (Figure 4).

**Figure 4:**
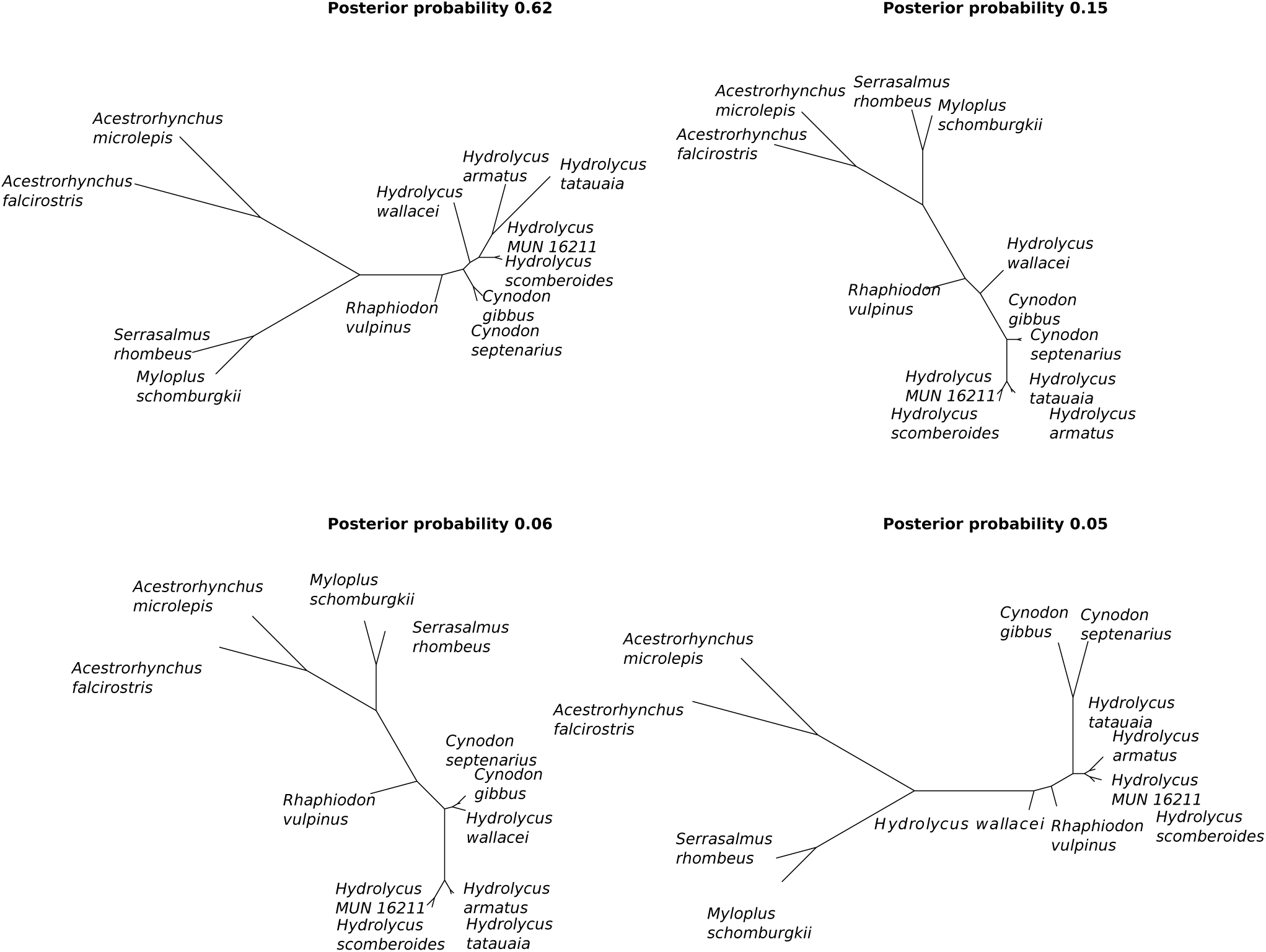
Summary of a posterior tree sample from the standard phylogenetic tree inference of the Cynodontidae dataset using MrBayes, which comprise nearly 90% of the tree posterior sample. These are labelled by posterior probability and sorted in decreasing order.

#### 3.4.2 Prior-posterior comparisons

It is important to compare the prior and the posterior in Bayesian analysis. The reason is that by comparing both sets of distributions we can identify cases where the information content in the likelihood is low, and therefore the posterior is being driven by the prior. This is particularly so for continuous parameters where we can plot the distribution of parameter samples as a kernel density or empirical density. Then we can compare the prior and posterior densities for a given parameter. Please note that we deactivate the data by rising the likelihood function to a power of 0 so that it is 1.0 in every MCMC iteration and thus sample just from from the prior densities. This is made by setting the option sampleFromPrior in beauti, or using the option underPrior=TRUE in the method mcmc.run of RevBayes. Then we will run another analysis for which we will have samples from the prior densities to compare with the full analysis which samples from the posterior. We should have now two trace or log files with parameter values.

The package tbea provides graphical methods for comparing prior and posterior densities (see the vignette prior posterior comparisons). For instance, the cross-plot can be used with a pair of log files, one for the prior (the *x*-axis) and one for the posterior (the *y*-axis) of the node ages in a divergence time analysis, so that the *x, y* pairs can represent prior and posterior node age means or medians, and error bars represent the 95% highest posterior density interval (Figure 5A). This kind of plot has been used in the literature when comparing prior and posterior MCMC samples, as well as when comparing the same kind of estimates coming from different independent runs or types of analysis (e.g., Drummond and Stadler, 2016; Álvarez-Carretero et al., 2022). This plot allows to see clearly that the prior node densities have more variance than the posterior ones.

**Figure 5:**
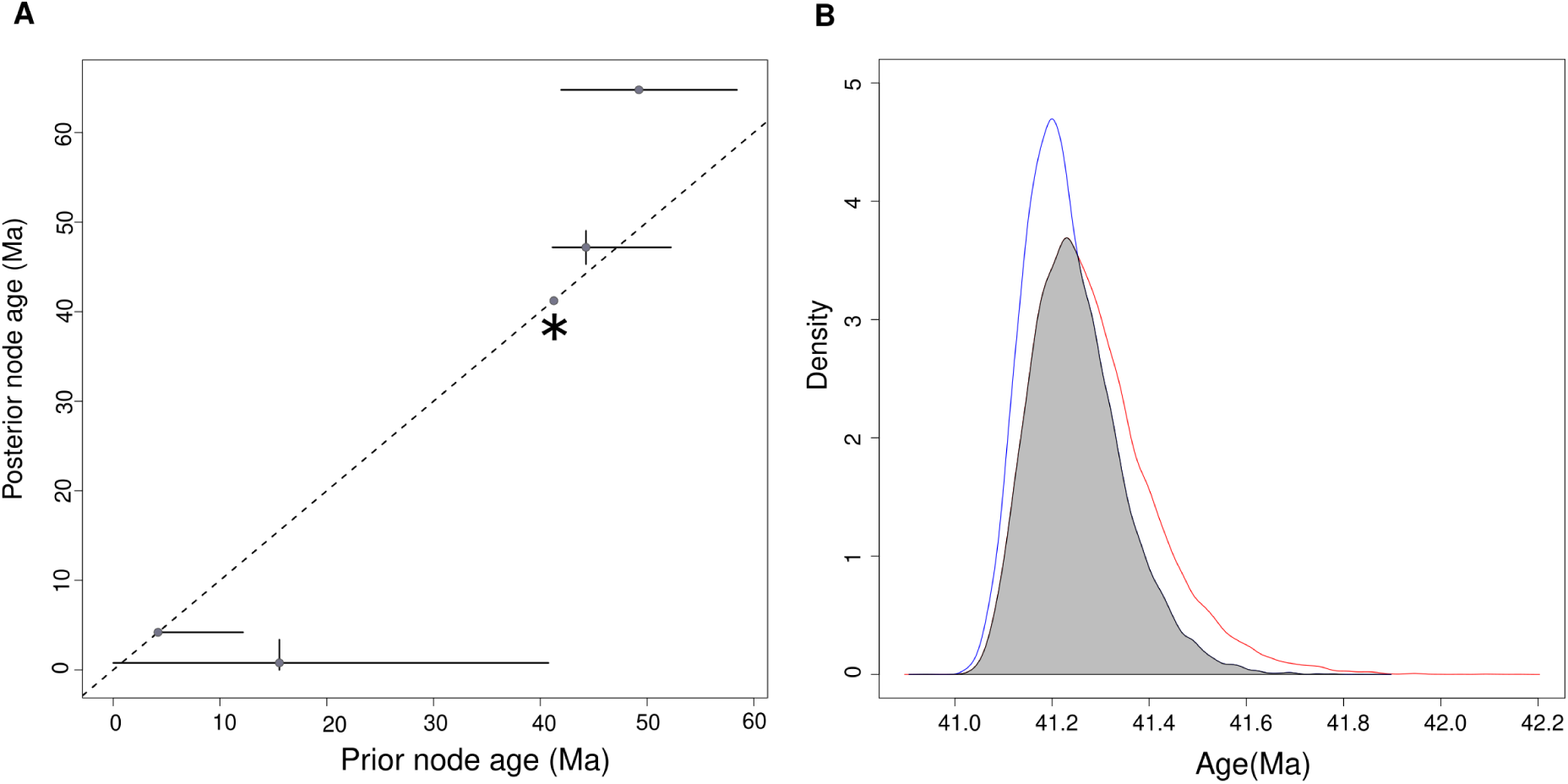
A. Cross-plot with a comparison between the prior and the posterior of node ages in the divergence time estimation of the Cynodontidae dataset. Note that he prior node densities have more variance than the posterior ones, which is a consequence of using the data through the likelihood function. B. Similarity between the prior (red) and posterior (blue) densities of the node *Hydrolycus*. The similarity, which is the value of the intersection (grey) between both distributions is 0.83.

Although visual comparison is useful and advisable, is some situations we may be interested in actually measuring the similarity between two densities, such as when comparing different calibration densities for the same analysis. We may also want to represent the degree of similarity between prior and posterior densities for a given parameter quantitatively. We can define similarity between two densities as the area under the intersection of both density curves. The function measureSimil achieves this and thus provides a descriptive measure of how similar two distributions are. The function can both plot the resulting distributions and their intersection (Figure 5B) as well as print out its value, or skip the plot and just return the value. When comparing the prior and posterior densities for a calibrated node depicting the genus *Hydrolycus*, we see that their intersection is quite large, 0.83, which may represent strong sensitivity of the posterior to the prior, and thus little impact of the likelihood. This strong sensitivity of the posterior is expected from analyses with few calibration points (Brown and Smith, 2018).

### 3.5 Inference of origination time given a set of time estimates

Let *X* be an arbitrary biogeographic area in geologic time bearing a biota *B_X_* . At a given point in time physical earth processes divide the area in two daughter areas *Y* and *Z* in a process commonly known as vicariance

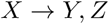

The expectation is that the biota *B_X_* also separates and reacts to this interruption of gene flow by speciation and thus creates a pattern of repeated sister-species-pairs between areas *Y* and *Z* (Figure 6A), although it might also undergo extinction at a given rate, but for simplicity we will assume first that there is no extinction taking place, so that

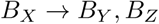

**Figure 6:**
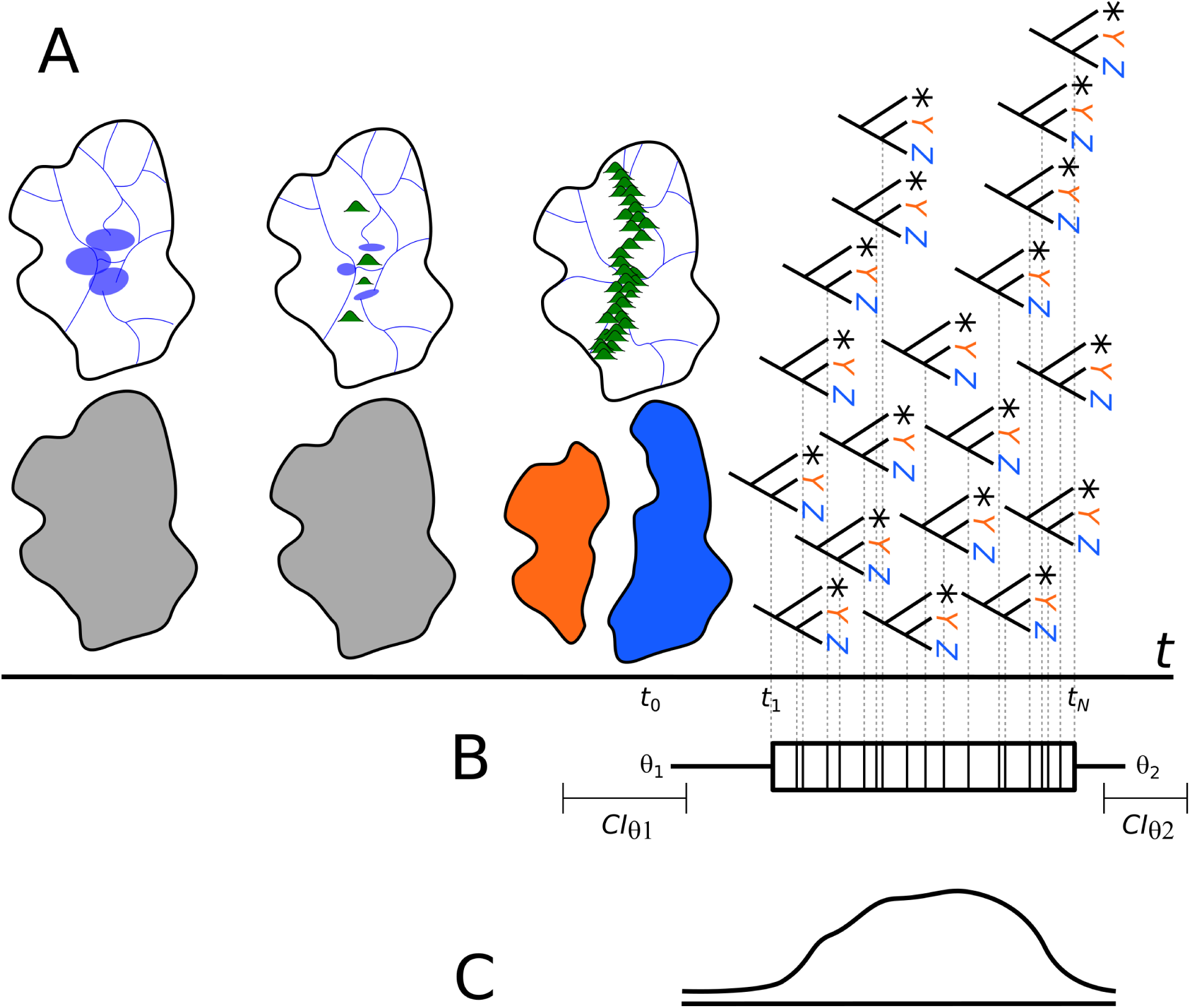
The inferential model and two possible analytical strategies depicted. A) The separation of an arbitrary biogeographic area *X* along geologic time that produces daughter areas *Y* and *Z*; after a lag in time after vicariance, the biota responds speciating and the collection of sister pairs distributed in daughter areas inform about the original time of separation *t*_0_. B) The specification of a stratigraphic interval and with endpoint parameters *θ*_1_*, θ*_2_, and all the occurrences in time *t*_1:*N*_ ; with such information it is possible to construct confidence intervals at a nominal *α* value for said parameters, in particular *θ*_1_. C) The empirical density of the divergence time events that allow to re-characterize it as a CDF and use regression for estimation of the *x*-intercept.

Note that the pattern preserved may also be subject to two obscuring processes: lack of speciation, and immigration from adjacent areas followed or not by speciation, thus creating non-*Y, Z* geographic patterns. We will again ignore these forces for sake of simplicity. Note that non-*Y, Z* patterns are uninformative on the process because they may happen before, coeval, or after the event *X* → *Y, Z*.

Now imagine the ancestral area *X* previously consisting of continuous lowlands with a common drainage network, now undergoes vicariance, for instance through mountain uplift, thus imposing a physical barrier to drainages that creates two different watersheds, and also, creates montane habitats. In sum, these two phenomena block gene flow in aquatic organisms, and promote speciation given that the lowland terrestrial biota is unable to get across mountain regions mostly due to physiological constrains. As the process also cuts the former drainage network creating two or more drainages that lack connectivity, the freshwater biota also looses gene flow, and respond speciating. This process is however not instantaneous in time, as both the geological processes will be in the scale of millions of years, and their response in the biota will also take some time after completion of area separation. This last phenomenon can be thought of as a lag between actual area separation *t*_0_ and the first speciation event *t*_1_ that will depend on how fast a particular biological group will respond to blocking of gene flow. Then, a number of speciation events in time *t*_1:*N*_ later than the true area separation *t*_0_ will take place as a consequence of the vicariance event (Figure 6A). The vector of speciation events is implicitly assumed to be measured without error because the probability that either endpoint parameter is actually present within the observed interval is zero following the definition of the confidence intervals as expansions beyond the first and last observed occurrences in time (cf. Equations 2,3). Given this inferential issue, outlier removal procedures might be necessary in order to remove suspicious data that are contradicted by the overall distribution of data (e.g., a divergence estimate that is too old in geologic time, or based on misleading calibration information).

As we lack direct evidence of the event *X* → *Y, Z* we aim at using information preserved in the statistical distribution of events *t*_1:*N*_ in order to estimate the time at which the vicariance event took place, *t*_0_ in the notation above. Herein we propose two alternative ways of looking at the statistical structure of these phenomena in time: First, as a stratigraphic interval formed by the unobserved origination of the vicariance event and its end (maybe unnecessary), along with the collection of events in time that resulted from it (Figure 6B, and second, as the distribution of these events in time (as well as its cumulative distribution) (Figure 6C).

Different estimates of the same parameter are often of interest in meta-analyses as well as means of specifying secondary node calibrations. However, how to summarise such information is not straight-forward (Hill, 2011; Hill and Miller, 2011). Here, we implemented three methods in which estimates from different distributions measuring the same parameter can be summarised in a single distribution: (i) using stratigraphic intervals (see vignette stratigraphic intervals), (ii) x-intercept confidence intervals (see vignette x intercept), and (iii) conflation of distributions (see vignette conflation). As an example, we estimated the age of the separation of drainages East and West of the Andes by using the first two approaches.

#### 3.5.1 Stratigraphic intervals

The vicariant model presented earlier shows an interesting distribution of events in time: They start with the origination time, and then happen for some time until the last component of the biota responds to the separation event. This pattern of events in time can be described by a stratigraphic interval. This is a collection of events in time for which the unobserved origination and extinction times are unknown. All the information we have comes from the set of occurrences in time (Figure 7). These models have been extensively used in palaeobiology and some flexible implementations have become recently available (Ballen, 2025). Although defined for species in the fossil record, we propose extending their meaning to something more general, such as sets of occurrences in time. This also includes their original applications to sets of time occurrences for a given species but also encompass sets of biogeographic occurrences such as vicariance events preserved in the phylogenies of multiple groups and triggered by a palaeogeographic event, for instance, drainage separation due to orogeny. It is important to note that nothing in the mathematical formulation of stratigraphic interval models is unique to species and therefore its application can be expanded without loss of correctness. The only difference in interpretation is that preservation parameters are now describing the distribution pattern of events in time.

**Figure 7:**
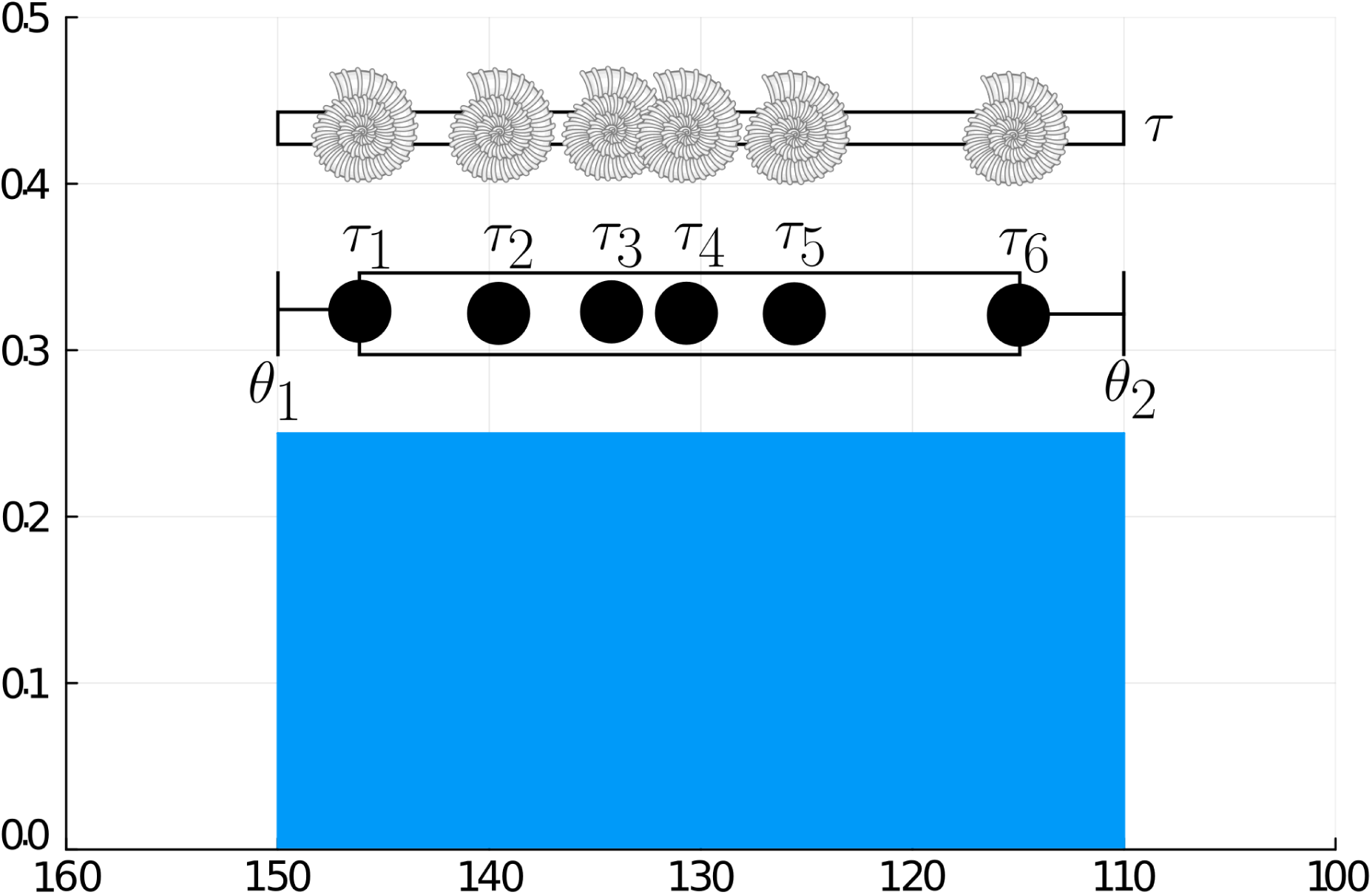
Stratigraphic interval. Suppose we have an Ammonite species through time. The species originates at time *θ*_1_ and becomes extinct at time *θ*_2_. Along this interval, it leaves preserved occurrences at times *τ*_1_*, . . . , τ*_6_. The underlying distribution is Uniform, which is governed by the preservation potential, and therefore the realisations of the preservation process. We estimate the origination and extinction times, as well as the preservation parameters from the data. Horizontal axis represents time from present (in Ma) and vertical axis represents density in the underlying distribution.

Frequentist approaches provide ways to estimate the confidence intervals on the endpoints of a stratigraphic interval, i.e., an estimate for the point of appearance and disappearance of a given taxon in geologic time. These methods are relevant here since we can think of a biogeographic events in the same way as taxa appear and become extinct. This approach is so far inexistent in historical biogeography and seems to fit the main goal of determining the final point of connection between biogeographic areas. Marshall (2010) summarised different approaches for estimating confidence intervals under certain sampling regimes, assumptions and confidence levels, ranging from classical to Bayesian estimators. The package tbea implements two of these methods, the constant-preservation estimator by Strauss and Sadler (1989) and the distribution-free estimator of Marshall (1994).

The first method provides estimators where one- (*θ*_1_) or two-parameter (*θ*_1_*, θ*_2_) cases are constructed adding and subtracting a portion *α* of the magnitude of the observed stratigraphic interval [*y, z*]. Such constant depends on the confidence level wanted (e.g., 95%).

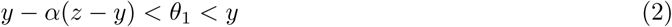

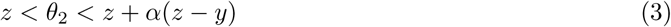

The constant *α* can be calculated either exactly (one-parameter case) or iteratively (two-parameter case).

The method of distribution-free confidence intervals was developed without assuming any particular distribution of underlying gap sizes, instead, it calculates quantiles of gap size between occurrence points for an ordered vector of gap sizes. The major cost of relaxing the assumption of a parametric approach in comparison with the constant-preservation method is that such intervals are larger, whereas the distribution-free approach require much more data for constructing intervals of the same width as the former (e.g., 95%).

The method works by constructing a confidence interval that has a level of confidence *C* (i.e., *Cth* percentile) for a *γ* confidence probability with the information from *N* gaps.

The lower bound is then the *i_th_* gap in the ordered vector applied to *min*(*vector*) − *x_i_* where *x_i_* is the magnitude of such *i_th_* element in the vector:

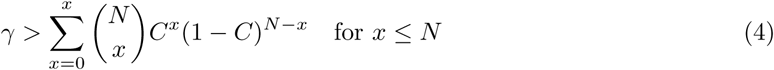

The upper limit can be constructed with the formula:

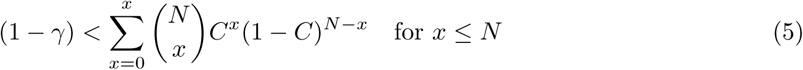

The method makes use of the binomial distribution in order to find the quantiles that satisfy the condition of being smaller that *γ* and from these picking the largest (in the case of the lower bound) and the smallest (for the upper bound). It is noteworthy that if only one *x* satisfies Equations (4) or (5), no lower or upper confidence interval can be constructed. Also, if *x* = *N* in (5), no upper confidence interval can be constructed since there is no (*x* + 1)*_th_* gap in the vector. Marshall (1994) mentioned that inference using his method is only valid as long as there is no trend in gap size (i.e., a trend of increase or decrease in gap size as we go along the time data). It is possible to use the Mann-Kendall test for monotonic trend to test such assumption (Mann, 1945; Kendall, 1948). Because the method works with quantiles of the sorted gap sizes, these are first calculated from the sorted vector of times, but then internally sorted when using the method “Marshall94”; however, whether there is any monotonic trend must be assessed on the unsorted version. We use the Mann-Kendall test on the times vector prior to applying the function.

These two methods are available in the function stratCI which allows to pick one of two methods: “straussSadler89” or “marshall94”.

As Marshall (1994) already noted, the method requires a large number of gaps to calculate both bounds for the 95% confidence interval; for our dataset with just 42 gaps (i.e., 43 times), we can find both bounds at the 80% confidence level (or lower), and thus claim that the lower and upper bounds for the 95% confidence interval are 7.07–7.2Ma. When we raise the confidence level to 95%, we can only infer the lower bound, thus being able just to claim that the 95% confidence interval is at least as old as 7.14 Ma. The higher the confidence level for finding the bounds on the 95% confidence interval, the better, in a similar way as calculating a 95% confidence interval is better than calculating a 70% confidence interval.

#### 3.5.2 x-intercept inference

Let us go back to the vicariance model presented earlier. One consequence of the separation event is that the biota does not necessarily respond immediately but instead with some lag depending on the lability to interruption of flow imposed by the separation event. For instance, in drainage separation via orogeny we expect that aquatic organisms will respond faster than terrestrial and volant ones. There is always some lag, small or not, of the accumulation pattern of vicariant events in time.

These separation events can be described as a distribution regardless of its generator process, which will follow a probability density function (PDF) *f* (*t*), with an associated cumulative density function (CDF) *P* (*X < x*). We can fit an empirical CDF and then use a linear regression model. The only case where a CDF would be truly linear would be in the case of a Uniform distribution where

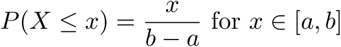

which implies that the empirical data in time follow a Uniform distribution. For any other case, a fitted CDF would be non-linear, and thus a linearisation should be applied to it. However, it is also possible to use robust non-parametric regression in order to avoid the issues with non-linearity of the CDF. Such regression model with an additional parameter *x*-intercept may be thought of as an estimator for the initial time *t*_0_ given the trend present in the linear model (Figure 8).

**Figure 8:**
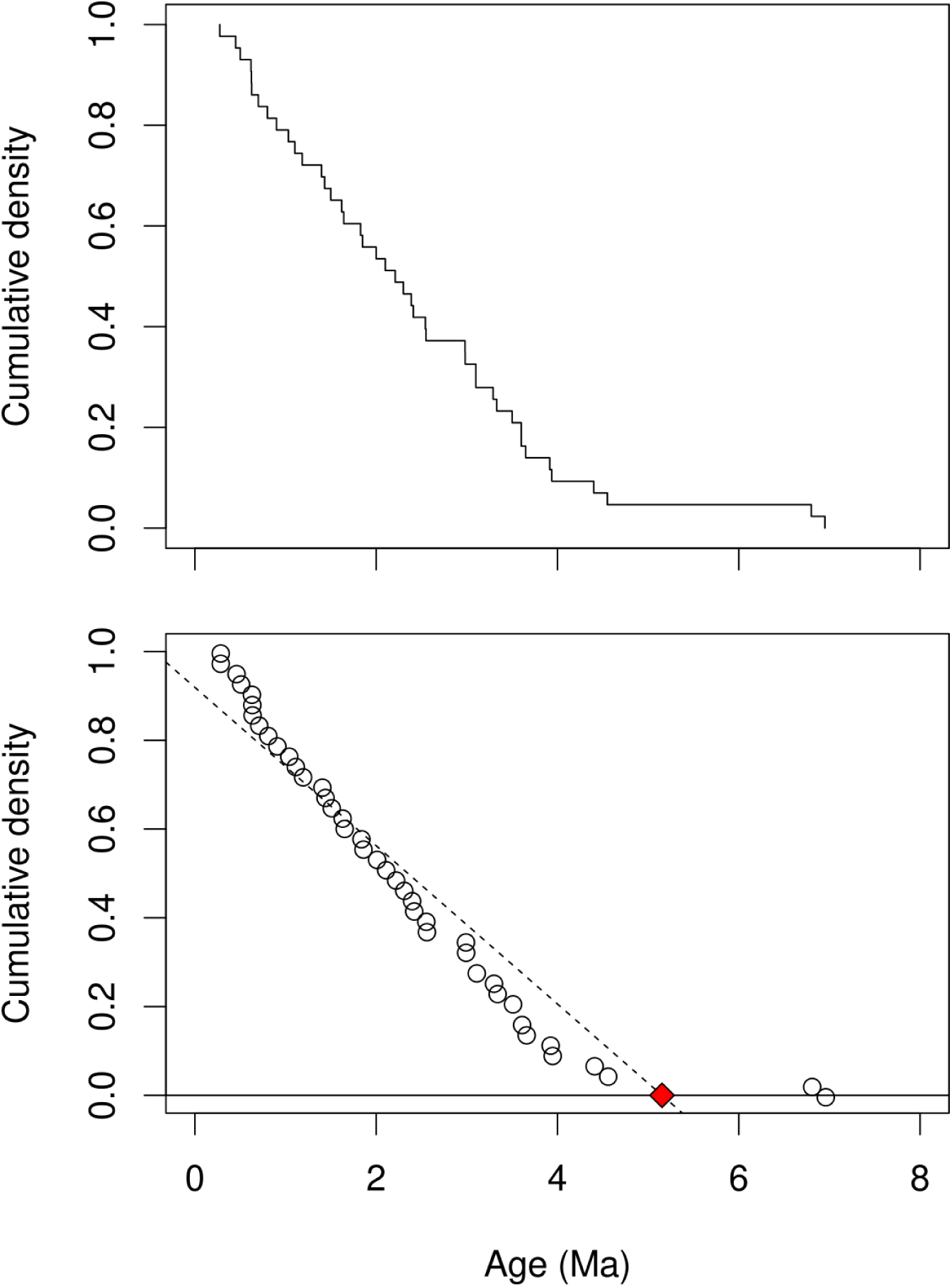
Empirical cumulative distribution of divergence time events. Above, plot of the empirical density; below, model fit of the cumulative distribution. The interrupted line represents the regression model on the cumulative values, and the red diamond represents the *x*-intercept.

Estimation of uncertainty regions such as confidence intervals on the *x*-intercept can be even more useful than the point estimate itself. Such regions can be constructed by several methods, for instance using bootstrap or analytical formulae for the predictive region or confidence intervals.

The linear model has the form

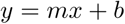

where *m* is the slope and *b* is the *y*-intercept. However, we are not interested in any of these parameters but instead in a third one, the *x*-intercept, that is defined as the point in *x* where *y* = 0. This parameter can be defined by letting *y* = 0 in the equation and thus

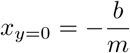

allows us to find the point at which the data began to be generated.

Several methods have been proposed for estimating the uncertainty on the *x*-intercept estimate, from the intuitive inverse regression *X* ∼ *Y* to bootstraping methods. According to Seber and Lee (2013, pp. 146-147) this problem has long been discussed in the statistical literature without much consensus, although some are of interest due to theoretical appeal, ease of implementation, or simplification of assumptions. Uncertainty on the estimate of the *x*-intercept can be obtained through Taylor series simplification for symmetric intervals. Another approach by Draper and Smith (1998, pp.83-86) allows the calculation asymmetric confidence intervals by projecting the values of the confidence region around the linear model on the *x*-axis by manipulation of the curves describing the confidence region, what gives

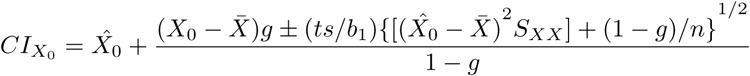

where *g* = *t*^2^*s*^2^*/*(*b*^2^*/S_XX_* ), *b*_1_ is the slope, *S_XX_* is the sum of squares of *X*, *t* is the *t*-statistic with n-2 degrees of freedom and at 1 − *α/*2 significance level, *s*^2^ is the standard deviation, X^0 is the estimationof the *x*-intercept, and ̄*X* is the mean of *X*. Here we need to fit a linear model in order to use the curves for the confidence region, and we can use classical least square models or robust non-parametric regression (Kloke and McKean, 2014).

It is also possible to use bootstrapping in order to estimate the variance and thus the confidence interval in asymmetric cases such as this one. Instead of using the estimates from a single fitted model on the coordinates of the empirical CDF, this method fits multiple models with random subsamples in order to construct a set of estimated values the *x*-intercept and this provides a confidence interval. The package tbea implements estimation of the confidence interval for the *x*−intercept using the method of Draper and Smith (1998) as well as estimation via bootstrap (see vignette x intercept).

Both require a linear model which can be set to one of the following: either ordinary least squares, or a robust linear model (Kloke and McKean, 2014).

Both methods produce similar results (Figure 9B). Although the method based on robust regression produced a CI wider than the one based on ordinary least squares, the bootstrap CI estimate was as wide as the two former combined. The fact that the bootstraping based on robust estimates is of more general application, it seems more adequate for these situations. It is noteworthy that these inferences are based on the assumption that the events in time are measured without error. However, this assumption relaxes due to the effect of re-sampling and uncertainty in the estimates. According to both estimators, the separation between cis- and trans-Andean areas took place between ca. 4.3 − 5.8 Ma.

**Figure 9:**
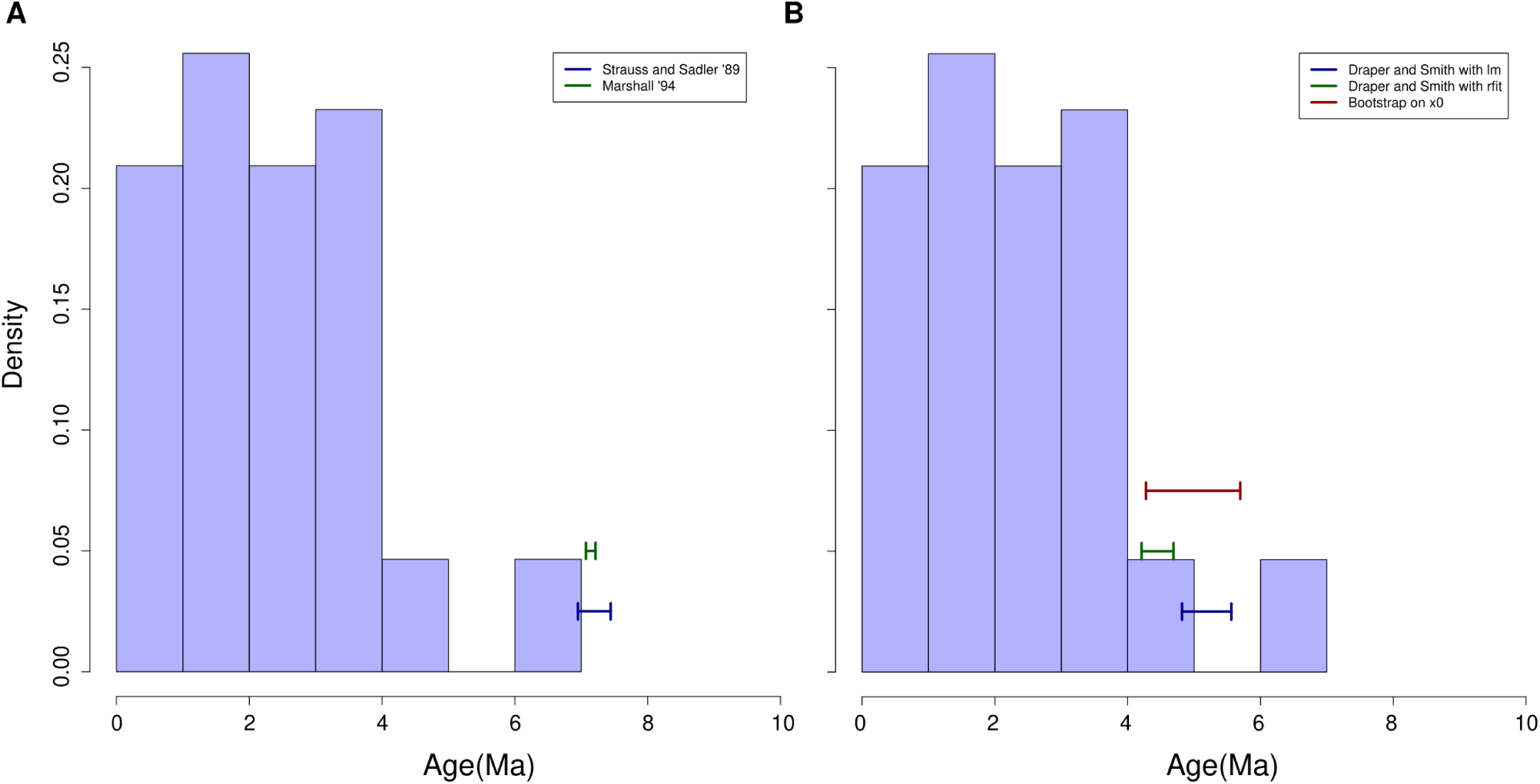
Estimation of the time of separation between cis- and trans-Andean drainages. A. Based on the methods of stratigraphic intervals of Strauss and Sadler (1989) and Marshall (1994). B. Based on the methods of CDF fitting using the estimator of Draper and Smith (1998) using two different approaches at model fitting, one using least squares traditional regression, and the other using robust regression (Kloke and McKean, 2014) and bootstraping techniques. Note that confidence intervals are near but no bound to be older than the oldest occurrence in methods based on CDF fitting, whereas they are forced to be older than the oldest occurrence in estimates based on stratigraphic intervals. This property of both methods can help to decide which is more reasonable to apply in specific circumstances, e.g., when the ages of the oldest occurrences are not very reliable.

These methods have produced results in line with those based on stratigraphic intervals. However, they allow estimate *x*-intercepts and consequently their confidence intervals that are not necessarily older than the oldest divergence time estimation event. Thus, they are robust to the assumption of measurement without error that is implicit in stratigraphic interval methods. Nevertheless, they are still sensitive to the presence of strong outliers as preliminary analyses showed that the regressions on these data with outliers affected strongly the robust regressions.

#### 3.5.3 Conflation of distributions

Combining different distributions into a single one is a useful tool for summarising different sources of information. Given a set of *N* distributions describing the same parameter, we aim at building a general distribution which incorporates information from the individual components. Multiple alternatives are available, and as with *x*-intercept estimation, the problem of summarising a set of distributions has several possible answers (Genest and Zidek, 1986). Hill (2011) discussed these methods and proposed the conflation as a better and general way of combining distributions describing the same parameter, and introduced the unary operator & as representing conflation. Let *Q* = &(*P*_1_*, . . . , P_N_* ) be the conflation of *N* densities describing the same time event *τ* :

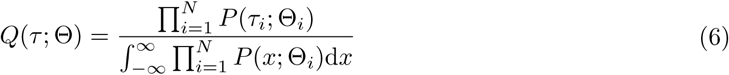

Among the useful properties of conflation we have that *Q* is also a distribution, which can be used for calculating probabilities. Also, the conflation combines the variance of each distribution, so that intuitively densities with more uncertainty (i.e., higher variance) will have a lower impact on the conflated distribution than others with lower variance, thus weighting the informativeness of all individual distributions appropriately.

Conflation proves useful when building secondary calibrations where more than two densities available for the same time event. Suppose that we have a biogeographic event with time *τ* , for which three different studies provide three independent densities, for instance, posterior probabilities for a node which corresponds to the biogeographic event. The best summary of these independent pieces of information is a conflation *Q*(*τ* ) = &(*P*_1_(*τ* )*, P*_2_(*τ* )*, P*_3_(*τ* )), which can be used as a secondary calibration point for a new divergence time estimation analysis as it is itself a PDF. Suppose that *P*_1_ = *N* (10, 1), *P*_2_ = *N* (13, 0.5), and *P*_3_ = *N* (14.4, 1.7). We can calculate the conflation of these three distributions.

Here, three Normals were used to approximate the posterior distribution of a given divergence time (Figure 10), and their conflation is depicted as a black line. It is evident that the conflated distribution shows more resemblance with the component with lowest variance (blue distribution), although incorporating the information coming from all of them. This distribution can in turn be used as a secondary calibration point in a subsequent divergence time estimation study, where a single node calibration must be used for a given node.

**Figure 10:**
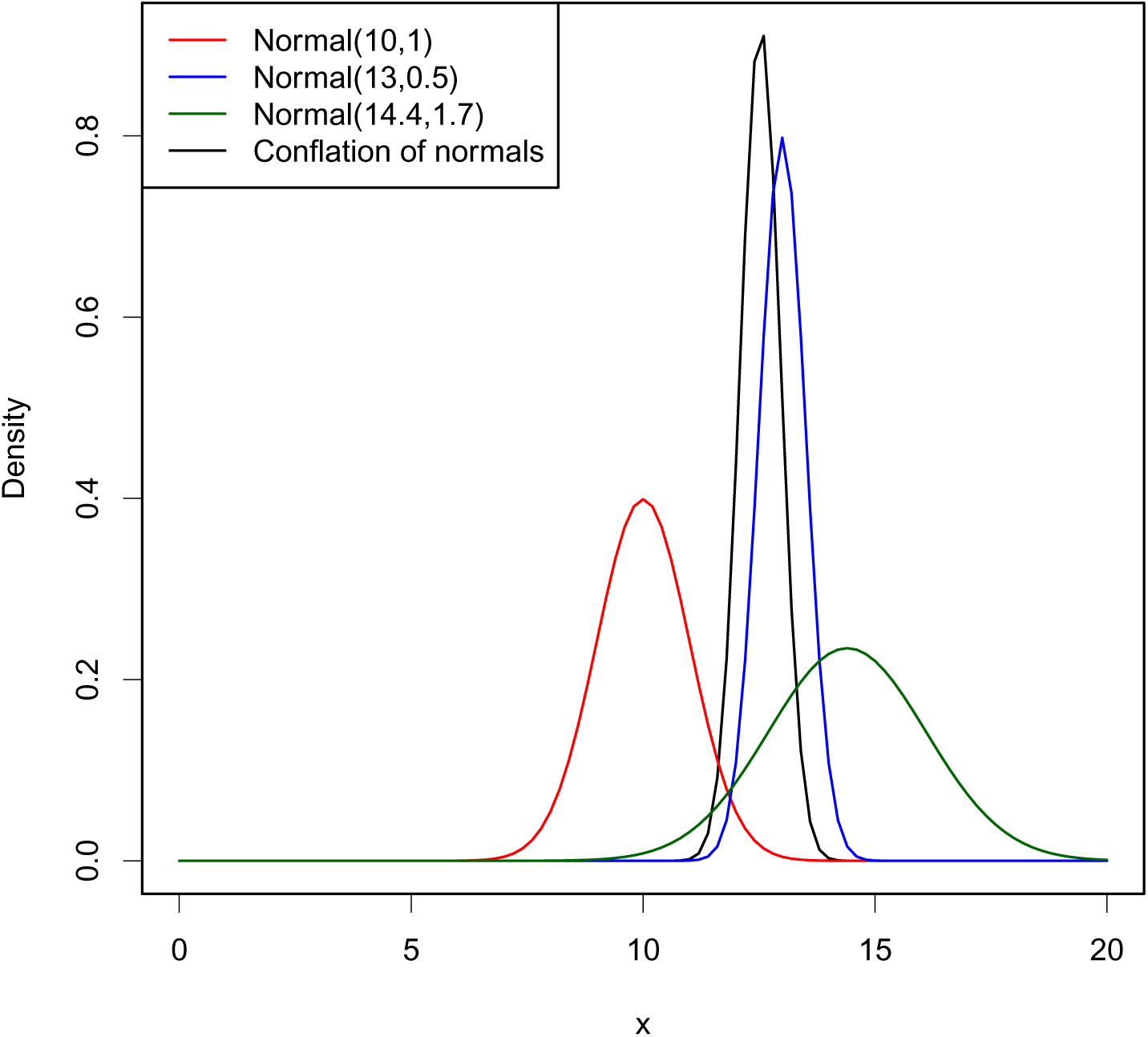
Conflation of three distributions representing imaginary posterior densities approximated by Normals of the same divergence time in Ma (red, blue, and dark green lines). The conflated distribution of these three represents the information coming from each proportional to their variance (black line).

The function conflate is used for building a conflation given arbitrary PDFs available in, that is, any function dDIST (e.g., dnorm, dexp, etc.). It uses a constructor function density fun that helps to specify the functions without need to calculate any object beforehand.

## 4 Discussion

Bayesian phylogenetic methods have had a dramatic impact in evolutionary biology due in part to its ability to integrate multiple sources of information and provide a natural way of characterizing uncertainty in parameter estimation. One of the challenges in empirical analyses is the reproducibility, although some programs have internal tools for improving it. Even though model specification is often carried out through the GUI program beauti, Beast2 will return the complete XML specification of the analysis as commented header of the log file containing the parameter values from the MCMC sampling. The downside of this strategy is that the XML specification is very challenging to understand by readers (although convenient from a development point of view). Both MrBayes and RevBayes support interactive sessions and the models can be specified piece by piece. However, their highest reproducibility standard is reached through the use of scripts, where the complete model is documented in code. MCMCTree is a special case where interactive mode is restricted to entering the path to the control file. The complete analysis needs to be specified in the control file and therefore, it qualifies as a script for practical purposes. The consequence is that any analysis specified in scripts is easier to understand and reproduce. Nonetheless, all these strategies are unable to document how the users chose the specific models, their parameters, and their distributions (e.g. priors). The pre-analysis tools in the present package aim at allowing the user not just find the best specification of distributions and model components, but also to document them as R scripts which can be shared alongside the analysis scripts. The same applies for post-analysis tools such as the comparison between model priors and posteriors, which are often verbally mentioned to have been done but not represented graphically in publications.

Techniques for estimating the origin time for a collection of events can be applied beyond the vicariant case herein presented. It is possible to use the same approach for a collection of events such as biotic interchanges (e.g., the Great American Biotic Interchange across the Panamà Isthmus) or colonisations (e.g., the invasion of South America by Gharials during the Cenozoic), which are in the end, collections of events happening in time but with a single origination time. Finally, the tools for estimating events in time from sets of point estimates, as well as the combination of different posterior densities from the same parameter are expected to improve the transparency of research choices when e.g. justifying the choice of secondary calibration points, or discussing the timing of biogeographic events when multiple sources are available for them. An additional advantage of this toolset is to integrate information from multiple areas of biological and earth sciences. They allow to make a better use of geological and palaeontological information for the specification of priors, as well as for estimating events in time, as some of the models (e.g. stratigraphic intervals) have been transferred here from palaeobiology to the analysis of Bayesian estimates. Distribution conflation or estimation of parameters in stratigraphic intervals using estimates from divergence time estimation analyses, in turn, can be helpful for research in palaeogeography and tectonics as they provide an independent source to contrast the predictions made with aid of geological tools. In the end, both areas are concerned with events in geologic time.

## Acknowledgements

This work was funded by FAPESP processes 2014/11558-5, 2016/02253-1, and 2023/07838-1 to GAB, 2020/13433-6 to Cĺaudio de Oliveira, and 2020/09442-0, 2020/10206-9 and 2022/01533-1 to SR. We thank the feedback by students at the Universidad de Antioquia in Colombia during a graduate course on phylogenetics in 2021, as well as the feedback from students in the graduate course Efficient Divergence Time Estimation using MCMCTree at Universidade Estadual Paulista in 2024, and help by Klauss Schliep when implementing metaprogramming code for numerical optimization. We thank both Uwe Ligges and Benjamin Altmann for their feedback during submission to CRAN which helped improve the documentation of the package. We thank T. Simõoes for feedback on functionalities of EvoPhylo. We thank the excellent and detailed feedback from the editor T. Guillerme and two anonymous reviewers which helped us improve the manuscript.

## Author’s contribution

Both authors contributed to conceptualization, development and writing.

## 5 Data availability statement

The R package tbea is hosted on CRAN (https://cran.r-project.org/package=tbea) and development versions are available on GitHub (https://github.com/gaballench/tbea). The stable version of the package can be installed with library(“tbea”) and the development version with remotes::install github(“gaballench/tbea”). The stable version herein described is v1.7.0, with DOI 10.32614/CRAN.package.tbea.

